# ORION: An agentic reasoning construct for the analysis of complex human immune profiling

**DOI:** 10.64898/2026.04.13.718286

**Authors:** Monica Dayao, Kenny Kim, Bernard Khor, Aaron Jaech, Bas van Opheusden, Aaron Bodansky, Joseph L. DeRisi

## Abstract

The capacity to generate high-dimensional biological datasets has outpaced the ability to interpret them. Technologies such as phage immunoprecipitation and sequencing (PhIP-seq) enable proteome-scale profiling of antibody repertoires, but interpreting thousands of enriched peptides into mechanistic hypotheses remains a labor-intensive bottleneck requiring expert synthesis of statistics, literature, and domain knowledge. Here we describe ORION (Omics Reasoning & Interpretation Orchestrator), a multi-agent framework that uses reasoning-capable large language models to perform end-to-end analysis of complex immune profiling data. ORION integrates statistical analysis, machine learning, and automated literature review into a single structured workflow, producing results that are reproducible and fully traceable. Applied to a published PhIP-seq dataset from autoimmune polyendocrine syndrome type 1 (APS-1), ORION recovered the canonical autoantibody signature in approximately two hours, closely recapitulating an analysis that originally required one to two months of manual effort. To test hypothesis-generation capacity on previously unseen data, we applied ORION to a novel PhIP-seq dataset from individuals with Down syndrome, for which no proteome-wide autoantibody reference exists. ORION distinguished disease from control samples with high accuracy, prioritized candidate autoantibody targets, and organized them into biologically coherent groups spanning immune, gut, and neuronal programs, generating testable hypotheses for experimental follow-up. These results demonstrate that agentic AI systems can compress the analysis of complex immune profiling data from weeks to hours, allowing scientists to redirect their time toward the fundamental biology.

## Introduction

The capacity to generate large and increasingly complex biological datasets continues to expand. Core technologies for interrogating biological systems are becoming faster, less expensive, and more capable, broadening the range of questions that can be addressed experimentally and computationally. Sequencing exemplifies this shift: whereas the first human genome required more than a decade and billions of dollars, modern high-throughput platforms can generate a human genome’s worth of data in a day for a few hundred dollars^1,2^. Similar reductions in cost and gains in performance across genomics, transcriptomics, proteomics, and additional modalities have driven a “multi-omics” era in which complementary, high-dimensional datasets are integrated to interrogate biological systems more comprehensively. In parallel, emerging approaches continue to expand what constitutes an “omic” dataset by enabling systematic measurement of biological processes that were previously difficult to assay at scale.

Phage immunoprecipitation and sequencing (PhIP-seq) is one such approach, enabling unbiased, proteome-scale profiling of antibody-antigen interactions^3–7^. By testing antibody reactivity against essentially all human proteins at high throughput, PhIP-seq has helped catalyze “autoreactomics”: the systematic study of autoantibody repertoires in health and disease^8^. However, unlike genomic, transcriptomic, or proteomic assays, PhIP-seq enrichment counts do not reflect simple molecular abundance; instead, they report antibody-peptide binding behavior shaped by affinity and avidity, epitope features, and immune history.

These properties impose distinct analytical challenges that are not readily addressed by standard “omics” pipelines. Autoreactivity is highly individualized, with no clear analogs to transcriptomic housekeeping genes or conserved baseline expression patterns. As a result, identifying biologically meaningful deviations—signals that rise above this background of individuality and may reflect shared autoimmune pathology—requires careful statistical modeling and context-aware interpretation. Individual experiments routinely yield thousands to tens of thousands of enriched peptides, and translating these enrichments into mechanistic hypotheses is time-consuming, often requiring expert synthesis of the literature and relevant databases.

Reasoning-capable large language models, such as GPT-5.2 and GPT-5.2 pro, have the potential to alleviate this bottleneck by automating early-stage statistical triage, machine-learning-based pattern finding, and literature/database review for high-dimensional data such as PhIP-seq. When integrated with an agentic reasoning pipeline and code execution environment, these systems prioritize antigenic targets, surface plausible biological connections, and generate testable hypotheses within hours. Here, we describe ORION, a multi-agent framework for analysis of ‘omic data. We present results with previously published data, as well as new data not available to the underlying model. These results suggest that as datasets grow and the atlas of human autoimmune disease profiles is populated, such AI-assisted analysis may enable faster, more consistent interpretation of new patient data and support increasingly nuanced, potentially predictive inference.

## Methods

ORION is a multi-agent framework for end-to-end PhIP-seq analysis organized around persisted intermediate outputs, auditable execution records, and iterative review (Figure 1). Starting from raw input data, it materializes a structured database, extracts cohort-aware features, runs supervised machine learning to prioritize candidate peptides and proteins, and retrieves supporting literature evidence – all orchestrated through a constrained agent loop that enforces reproducibility. All computational steps execute in a sandboxed Python environment.

**Figure 1.**
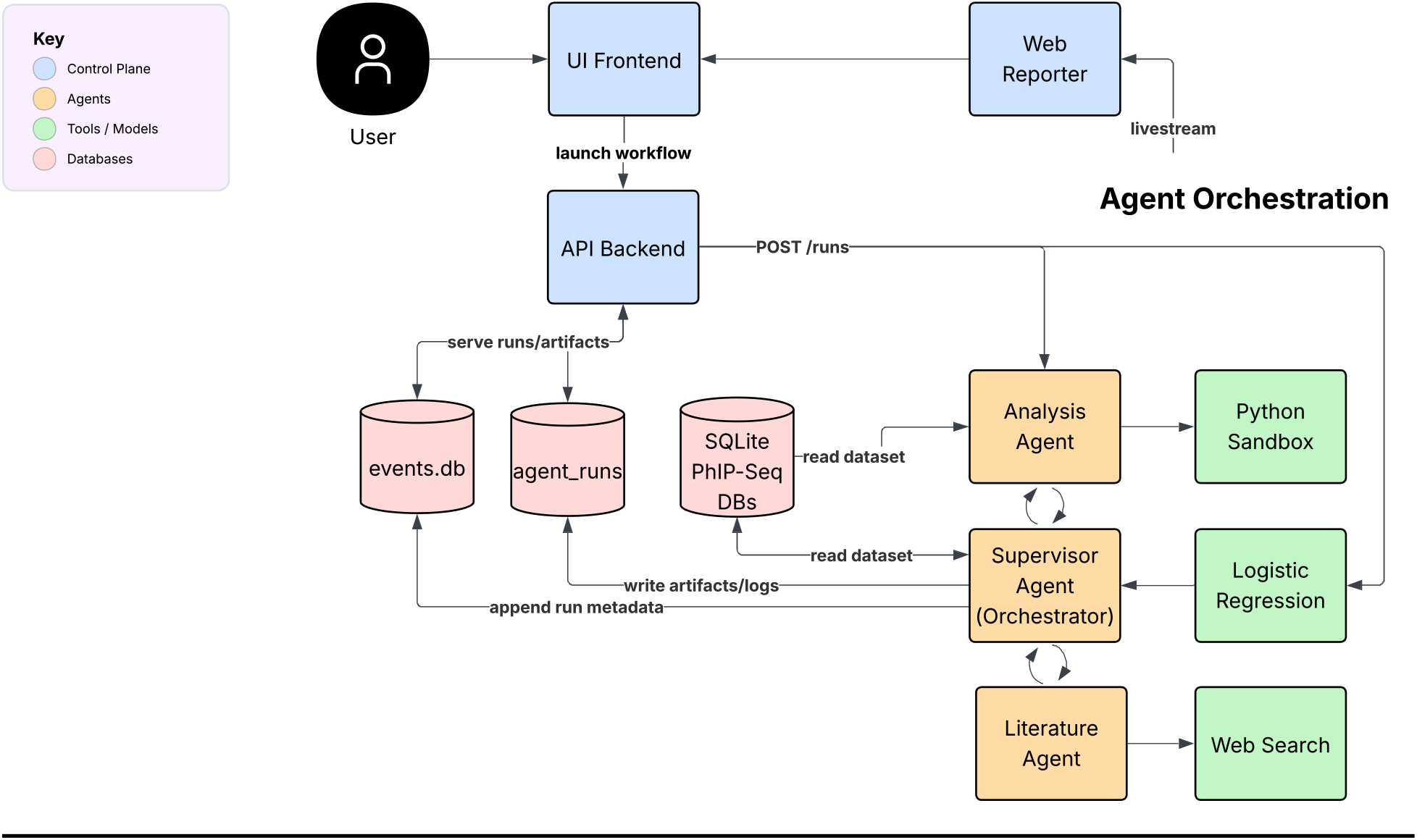
ORION multi-agent workflow architecture. ORION orchestrates end-to-end PhIP-seq analysis through three specialized agents (a Main Analysis Agent, a Literature Agent, and a Supervisor Agent/orchestrator) operating in a constrained Plan → Execute → Verify loop that enforces reproducibility and auditability through persisted, machine-readable intermediate outputs and execution logs. The Supervisor Agent monitors artifact completeness and triggers retries as needed, while a web reporter streams live progress to the user frontend. Run metadata and artifacts are made persistent via a SQLite database, and all computational steps execute within a sandboxed Python environment.

**Figure 1.**
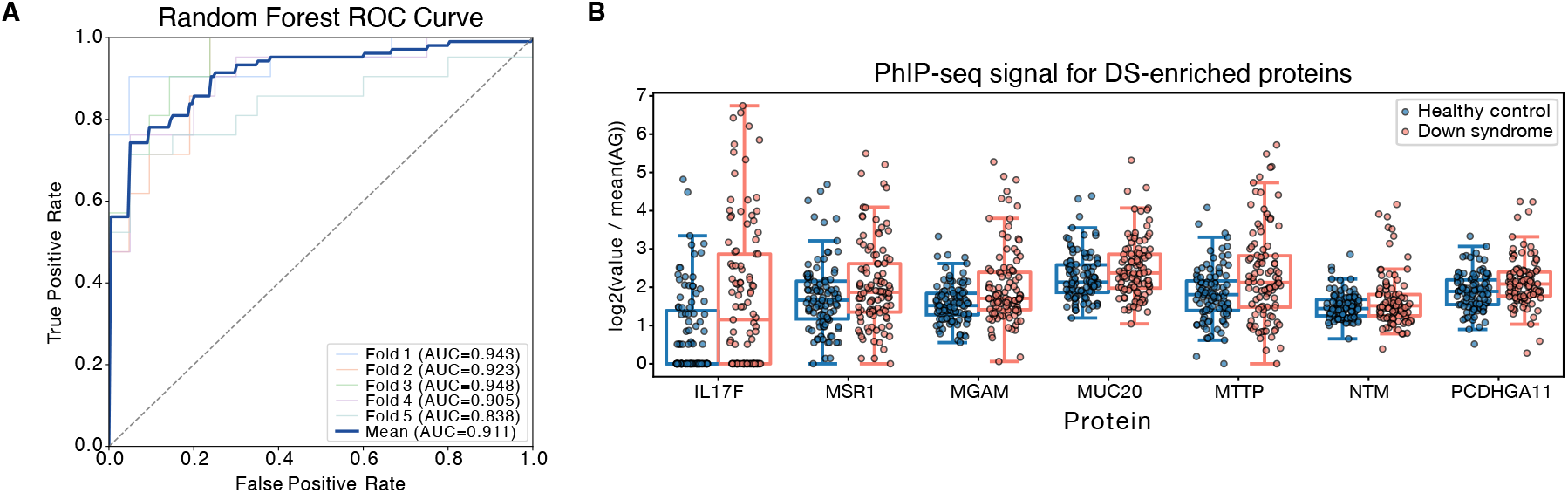
ORION-prioritized candidate proteins and classifier performance in Down syndrome versus healthy controls. A) Receiver operating characteristic (ROC) curves for the random forest model trained by ORION to discriminate between Down syndrome and healthy control samples, summarizing cross-validated performance (mean AUC = 0.911). B) Log2 AG-normalized RPK values are shown for a subset of lower-prevalence, higher-specificity candidate proteins prioritized by ORION, spanning immune-modulatory (IL17F, MSR1), gut/barrier-associated (MGAM, MUC20, MTTP), and neuro-adhesion-associated (NTM, PCDHGA11) programs. Each point represents an individual participant, stratified by group.

Importantly, ORION is a generalizable framework, appropriate for a large range of biologic ‘omics data. It preserves the core framework architecture and execution model described here. The complete ORION code and schema are available at github.com/openai/orion-multistep-analysis.

### Input Data

ORION accepts either (i) a paired metadata table and peptide enrichment matrix or (ii) a prebuilt SQLite database. The metadata table must include a sample identifier and label column; label aliases are normalized to canonical labels: D (disease), HC (healthy control), and AG (mock-IP AG-beads-only samples), while any other labels are preserved and uppercased. The peptide matrix is a wide sample × peptide table where the first column is sample ID and remaining columns are peptide identifiers.

### Dataset Materialization

Input matrices are converted into a sparse SQLite schema to reduce storage and accelerate feature extraction. At minimum, the database contains: samples (canonical sample IDs and labels), peptides (peptide identifiers), and rpk_sparse (only nonzero entries, sample × peptide → value). No statistical normalization is applied during import.

### Candidate Prioritization

ORION generates candidate peptide and protein lists using three complementary approaches:

- Cohort-aware peptide summaries derived from SQLite queries (prevalence, nonzero abundance summaries, and label-stratified contrasts).
- Supervised feature attribution via logistic regression (D vs HC; details below).
- Protein-level aggregation, when a peptide → protein mapping is available.

Prioritized candidates are subsequently passed to the Literature Agent for evidence retrieval and biological contextualization (see Agent Pipeline and Prompting).

### Logistic Regression Module

Only D and HC samples are used (minimum 2 per class); AG and any other labels are excluded. A dense sample × peptide matrix is constructed from rpk_sparse with missing pairs filled as 0.0, and peptides are filtered out if all labeled samples have RPK < 5.0 (pseudocounts=False); constant features (nunique ≤ 1) are also dropped, and no log transform or scaling is applied. A scikit-learn^9^ LogisticRegression model with L2 regularization (C = 0.1, solver = lbfgs, max_iter = 1000) is fit using StratifiedKFold cross-validation with n_folds = min(5, n_per_class). Coefficients are averaged across folds per peptide (mean_coef) and ranked by abs_coef = |mean_coef|, where the sign of mean_coef indicates enrichment direction. When a peptide → protein mapping is available, protein-level scores are computed as aggregate_abs_coef = ∑ |mean_coef_peptide| and sum_coef = ∑ mean_coef_peptide across all mapped peptides. By default, the top_k = 25 peptides and proteins are reported with full coefficient summaries.

- **Agent Pipeline and Prompting**. ORION orchestrates the above steps through three specialized agents operating in a constrained loop in which each stage produces persisted intermediate outputs and execution logs that make the analysis traceable and auditable. Each agent is governed by a structured system prompt that defines its role, required outputs, and operational constraints (full prompt texts are provided in Supplementary Materials).
- **Main Agent (Analysis Agent):** The prompt specifies a mandatory four-phase workflow: data sanity and setup, broad-scan differential analysis across ≥100 proteins/peptides, multi-method quantitative inference, and mechanistic interpretation including competing disease models. The prompt enforces AG-bead normalization consideration, prevalence-based framing of results, dual peptide- and protein-level reporting, and traceability via persisted SQLite output tables. It prohibits the agent from fabricating external knowledge and requires all external facts to be requested through the Literature Agent interface.
- **Literature Agent:** The prompt constrains the agent to return a single structured JSON object containing protein-level annotations (function, subcellular localization, tissue expression, pathway membership, disease associations, and immune-accessibility implications), cluster-level synthesis across multiple query proteins, null-evidence reporting when data are sparse, and scored relevance heuristics. It prioritizes primary databases (PubMed, UniProt, OMIM, Human Protein Atlas) over secondary sources and instructs the agent to focus on normal protein biology rather than restricting searches to autoantibody-specific literature.
- **Supervisor Agent:** The prompt implements a nine-item checklist (data sanity, AG normalization, multi-method analysis, broad-scan coverage, justified narrowing, mechanistic interpretation, competing models, literature compliance, and traceability) with explicit loop-prevention rules—if the Analysis Agent fails to satisfy an item for the same reason twice, it is marked as constrained and the supervisor proceeds rather than re-requesting. Termination requires all mandatory checklist items to be satisfied.

This Plan → Execute → Verify loop ensures that analysis steps are reproducible and that conclusions are traceable to specific computational outputs.

## Results

### APS-1 benchmark recapitulates a known autoreactive signature

We evaluated ORION on a published PhIP-seq dataset from autoimmune polyendocrine syndrome type 1 (APS-1), a monogenic disorder with a well-characterized autoantibody repertoire^4^. Starting from raw enrichment matrices, ORION executed artifact-aware preprocessing, candidate prioritization, and literature-integrated interpretation. The framework recovered the canonical APS-1 autoreactive signature: anti-cytokine antibodies (IL-17F, IL-22, and multiple IFN-α subtypes), steroidogenic enzyme autoantigens (CYP11A1; 52.2% disease vs. 0% controls), and organ-specific endotype markers including RFX6 (gastrointestinal), NLRP5 (parathyroid), SOX10 (melanocyte), and KHDC3L (ovarian insufficiency). A composite signature score across these targets separated disease from non-disease samples. Overall, these results closely recapitulated the published manually annotation analysis. Total runtime was 2 hours and 16 minutes. In contrast, the original manual analysis consumed approximately 1 to 2 months, conservatively.

### ORION identifies a PhIP-seq signature in Down syndrome after artifact-aware normalization

In the previous example, the dataset that was used as input to ORION was previously published, and thus available to the underlying reasoning model. To assess the utility of ORION for a dataset that has never been publicly disclosed, and for which no previous immunological signature has been described, we tasked ORION to analyze a PhIP-seq dataset derived from 105 individuals with Down syndrome (DS; trisomy 21), 103 age-matched healthy controls (HC), and 17 bead-only controls (AG), a control for non-specific phage binding to Protein A/G beads. The dataset comprised 563,322 peptides mapping to 19,949 proteins and was highly sparse (≈86% zeros). ORION completed the full analysis in 1 hour and 41 minutes.

To address stochastic, non-specific bead binding, ORION compared AG-normalized and unnormalized results. Without AG normalization, prevalence rankings were dominated by recurrent non-specific features, obscuring cohort differences. AG normalization reduced these artifacts and altered prevalence calls accordingly (e.g., MUC16 shifted from ∼76–77% positivity in both DS and HC to ∼0–1% after AG normalization). ORION therefore carried AG-normalized outputs forward for both binary prevalence calls and continuous enrichment scoring.

### DS–HC separation reflects distributed signal and resolves into multiple interpretable programs

Using the AG-normalized protein feature space, ORION quantified DS–HC differences using complementary statistical tests and supervised models with cross-validation. Classification supported a distributed signal: L1-regularized logistic regression achieved mean AUC = 0.769 and random forest achieved mean AUC = 0.911 (Figure 2A). In contrast to APS-1, DS separation was not driven by a small set of highly prevalent antigens, and feature attribution was distributed across partially overlapping signals.

Across methods, the most consistently DS-enriched proteins were predominantly intracellular, including RPUSD4 (28.6% DS vs 6.8% HC), BNIP1 (27.6% vs 7.8%), and PPIF (19.0% vs %). In addition, ORION identified multiple lower-prevalence, higher-specificity candidate programs enriched in subsets of DS participants and uncommon in HC (≤5% for most candidates). These included (Figure 2B):

1. **Immune-modulatory targets**: IL17F (8.6% DS vs 1.0% HC) and MSR1.
2. **Gut/barrier-associated targets**: MGAM and MUC20, with MTTP as a convergent enterocyte-associated marker.
3. **Neuro-adhesion–associated targets**: NTM and PCDHGA11, with APBB2 and FRMPD4 as convergent neuronal-context candidates.

### Ranked candidates motivate discriminating experimental next steps

ORION translated ranked features into testable experiments aimed at distinguishing among candidate mechanisms. Prioritized next steps included: correlating interferon-stimulated gene signatures and STAT1 pathway readouts with autoantibody breadth; relating MGAM/MUC20/MTTP reactivity to biomarkers of gastrointestinal inflammation and barrier disruption with orthogonal binding assays; and testing extracellular binding to NTM and PCDHGA11 using cell-based assays, with paired CSF testing where available.

## Discussion

ORION has immediate implications for the analysis of complex immune profiling data because it converts an often bespoke, expert-driven workflow explorations into a reproducible end-to-end process: cohort-aware statistical triage, supervised feature attribution, and literature-informed biological synthesis that yields explicit, tractable follow-up experiments. Run logs of the ORION Plan → Execute → Verify loop provide transparent and traceable records of each run. Unlike other high dimensionality biological data, like single cell sequencing, PhIP-seq is a stringent test case because its measurements are not simple abundance readouts; enrichment reflects antibody– peptide binding shaped by affinity/avidity, epitope features, and immune history. Moreover, unlike other ‘omics that benefit from shared reference features (e.g., housekeeping genes), each individual’s autoreactive repertoire is highly idiosyncratic, producing variable baselines. As a result, interpretation typically requires context-aware analysis coupled to substantial domain knowledge. ORION’s core innovation is to effectively integrate these steps into a single structured loop that can be applied consistently across large datasets, rather than relying on *ad hoc* analyses that vary by analyst and are constrained by time available. Although PhIP-seq is the demonstration modality used here, ORION is designed to be a generalizable framework that can be adapted to a broader range of ‘omic-driven biological data analysis.

A key practical consequence is efficiency. While statistical feature discovery can be automated, also integrating a fragmented literature into a coherent mechanistic model and a concrete experimental plan requires training and expert biological intuition. ORION compresses this end-to-end cycle so interpretation can plausibly keep pace with data generation, making large retrospective backlogs tractable for hypothesis generation, which in turn can be audited, reviewed, and refined by researchers before being put into action. In APS-1, where the autoantibody landscape is well defined, ORION recovered the expected autoreactivity signature and reproduced canonical separation from healthy controls, validating the end-to-end workflow. However, as noted above, the underlying model was exposed to APS-1 publications, and thus performance should be interpreted as evidence of coherent execution rather than de novo biological discovery.

The Down syndrome analysis is more informative as a hypothesis-generation test, because no proteome-wide autoantibody or PhIP-seq reference dataset exists in the literature, and the data used here are among the first of their kind. In this setting, ORION’s ability to integrate a broad body of published immunobiology becomes a practical advantage, enabling it to retrieve and synthesize relevant evidence at a scale that is time consuming and labor intensive to achieve manually. The results support a heterogeneous, interferon-aligned autoreactivity pattern consistent with trisomy-21 immune remodeling and the interferonopathy-like baseline state described in prior studies^10–13^. DS–HC separation was not driven by a small, highly shared autoantigen class; instead, it emerged from a distributed signature consistent with partially overlapping autoreactivity programs across individuals. Indeed, ORION’s classification task achieved an AUC of 0.91, which is remarkable for any human immune profiling-based assessment. Within this global context, ORION prioritized a smaller set of subset-enriched, immune-accessible candidates and organized them into plausible tissue-linked hypotheses— including barrier-associated and neuro-adhesion–associated programs—while preserving a clear boundary between statistical association and mechanistic inference. Together, this ranking, synthesis, and explicit treatment of uncertainty is intended to shorten the path from large lists of enriched peptides to a small number of coherent, discriminable models suitable for direct testing.

We also note the potential clinical relevance of applying ORION to pediatric immune-mediated disease. Down syndrome is among the most common genetic conditions associated with immune dysregulation, affects more than 200,000 individuals in the United States^14^, and confers substantially increased risk of severe infection, autoimmunity, and other immune-related complications beginning in childhood^15,16^. Yet the mechanisms underlying this altered immune state remain only partially defined. In this context, ORION’s ability to identify candidate autoreactive targets and organize them into coherent biological programs may be particularly valuable for pediatric disorders, where mechanistic insight is often limited despite substantial clinical burden. In the present dataset, for example, ORION prioritized autoantibodies targeting IL17F, a cytokine involved in mucosal immunity and host defense that lies within a therapeutically actionable immune signaling axis. If follow-up studies confirm that these antibodies are functionally relevant, such findings could clarify one contributor to immune dysregulation in Down syndrome and help nominate more precise therapeutic strategies. More broadly, many pediatric diseases with suspected immune dysregulation remain mechanistically unresolved despite the existence of high-dimensional immune-profiling data. By accelerating interpretation of such datasets, ORION may help convert clinically important but analytically underused data into experimentally tractable hypotheses across a wider range of pediatric disorders.

ORION also has important limitations. Reasoning-capable systems can produce explanations that are plausible but wrong, so interpretation must remain tightly grounded in the pipeline’s statistical outputs and treated as provisional until experimentally validated. As with any hypotheses, follow up suggestions will require scrutiny and refinement. Deployment also faces practical constraints: compute requirements and per-query cost can be substantial, especially for large cohorts and iterative exploration, and should be managed through configurable model selection, iteration limits, and adjustable analysis depth, with local execution options where appropriate. Finally, although PhIP-seq is a demanding demonstration, the broader goal is modality-agnostic. ORION is designed to extend to other ‘omics with appropriate adaptations to input schemas, feature extraction, and prompting, including proteomic and transcriptomic measurements and T cell/B cell repertoire sequencing.

As immune profiling studies grow in scale and become increasingly longitudinal, such systems could enable cross-cohort matching to identify individuals who share rare immune programs, and support per-person monitoring for earlier detection and prognostic inference in immune-mediated disease and cancer, especially with the integration of associated clinical meta-data. The central innovation is that ORION converts high-dimensional, unfamiliar immune profiling data into coherent, testable models at a speed and scale no human team can match, while preserving explicit uncertainty and auditability. These results suggest that early bottlenecks in data analysis of complex human ‘omics data can be rapidly overcome, shifting the bulk of expert scientist time toward fundamental biology: experimental design, experimental execution, model building, and hypothesis testing.

